# Skeletal Muscle Mitochondrial Morphology Negatively Affected by Loss of Xin

**DOI:** 10.1101/2023.09.14.557614

**Authors:** Grace Martin, Dhuha Al-Sajee, Molly Gingrich, Rimsha Chattha, Michael Akcan, Cynthia M.F. Monaco, Megan C. Hughes, Christopher G.R. Perry, Irena A. Rebalka, Mark A. Tarnopolsky, Thomas J. Hawke

## Abstract

Altered mitochondrial structure and function are implicated in the functional decline of skeletal muscle. Numerous cytoskeletal proteins have been reported to affect mitochondrial homeostasis, but this complex network is still being unraveled. Here, we investigated alterations to mitochondrial structure and function in mice lacking the cytoskeletal adapter protein, Xin.

Xin deficient (Xin-/-) and wild-type (WT) littermate mice were fed a chow or high-fat diet (HFD; 60% kcal fat) for 8 weeks before high-resolution respirometry, histology, electron microscopy and Western blot analyses of their skeletal muscles were conducted.

Immuno-electron microscopy and immunofluorescence staining indicates that Xin is present in the mitochondria and peri-mitochondrial areas, as well as the myoplasm. Intermyofibrillar mitochondria in chow-fed Xin-/- mice were notably different from WT; frequently spanning a whole sarcomere and/or swollen in appearance with abnormal cristae. Succinate Dehydrogenase and Cytochrome Oxidase IV (COX) activity staining indicated greater evidence of mitochondrial enzyme activity in Xin-/- mice. HFD did not result in a difference between cohorts with respect to body mass gains or glucose handling. However, electron microscopy revealed significantly greater mitochondrial density (∼2.1-fold) with evident structural abnormalities (swelling, reduced cristae density) in Xin-/- mice. Complex I and II-supported respiration were not different between groups per mg muscle, but when made relative to mitochondrial density, were significantly lower in Xin-/- muscles. Western blotting of fusion, fission, and autophagy proteins revealed no differences between groups.

These results provide the first evidence for a role of Xin in maintaining mitochondrial morphology and function but not in regulating mitochondrial dynamics.

## Introduction

Skeletal muscle is the largest organ in the human body and is principally responsible for both our physical and metabolic capacities. Accounting for up to 80% of post-prandial whole-body glucose uptake (Thiebaud et al., 1982; DeFronzo et al., 1985), skeletal muscle’s high capacity for fuel usage is necessary to meet the metabolic demands of this tissue. The metabolic capacity of skeletal muscle is largely established by the mitochondrial population, which comprises ∼4-7% of skeletal muscle volume (Lundby & Jacobs., 2016; Meinild Lundby et al., 2018). Mitochondria are essential for energy production and use oxygen and fuel substrates to generate energy in the form of adenosine triphosphate (ATP) for energy-dependant processes in the cell.

Impairments to mitochondrial structure or function can result in disrupted fuel handling and lead to the development of metabolic diseases such as, obesity and type 2 diabetes (Sivitz et al., 2010). Studies of skeletal muscle from individuals with type 2 diabetes have repeatedly shown a downregulation of genes involved in mitochondrial bioenergetics (Patti et al., 2003; Mootha et al., 2003), as well as the decreased expression and activity of related proteins (He et al., 2001; Kelley et al., 2002; Ritov et al., 2005). While not without some debate, it has also been proposed that mitochondrial dysfunction can lead to insulin resistance directly, with impairments to fat oxidation causing the accumulation of lipid metabolites and subsequent interference with the insulin signaling pathway (Kelley et al., 2002). Taken together though, it is generally well agreed that the maintenance of a healthy skeletal muscle mitochondrial pool is critical for the maintenance of whole-body metabolic function. Thus, it is of the utmost importance to identify proteins involved in mitochondrial homeostasis in order to identify novel players involved in mitochondrial dysfunction and new therapeutic strategies to improve the metabolic and physical capacities of skeletal muscle.

The interplay between proteins of the actin cytoskeleton and the intermyofibrillar mitochondria have been implicated in myofiber homeostasis (Rappaport et al., 1998; Appaix et al., 2003; O’Rourke et al., 2018). Xin binds and organizes the actin cytoskeleton with a predominant presence in striated muscles (Pacholsky et al., 2004; van der Ven et al., 2006). Though Xin was originally thought to be localized to the myotendinous junction (MTJ) within skeletal muscle (Feng et al., 2013), we and others have since reported Xin within resting skeletal muscles and their progenitor cell population (i.e. satellite cells), as well as a robust upregulation of Xin in response to muscle damage in both humans and rodent models (Hawke et al., 2007; Otten et al., 2012; Nilsson et al., 2013; Orfanos et al., 2016). Histologically, mice with a whole body knockout of Xin (Xin-/-) display in a mild myopathic phenotype, but functionally exhibit a significant increase in muscle fatigability and greatly reduced force recovery (Al-Sajee et al., 2015). To identify a mechanism underlying the impaired muscle function testing in Xin-/- mice, and given the aforementioned importance of cytoskeletal proteins to mitochondrial function, we undertook an investigation of skeletal muscle mitochondria in mice deficient for Xin and their WT littermates.

## Methods

### Animals

All animal experiments conducted were approved by the McMaster University Animal Care Committee and were in accordance with the Canadian Council for Animal Care Guidelines. C57Bl/6 mice heterozygote for the full body deletion of Xin (the gene product of Xirp1; Xin+/-) were a generous gift from Dr. Dieter Fürst, University of Bonn, Germany; Otten et al., 2010). These mice were bred to create Xin deficient (Xin -/-) and wild-type (WT) littermate mice that were used in the following studies. All animals were housed under controlled conditions of 21°C, 50% humidity and 12 hour/12 hour light-dark cycle. Animals were provided enrichment materials and had ad libitum access to food and water. All mice were fed a standard chow diet [Research Diets (New Brunswick, NJ), D1245K Rodent Diet: energy (kcal/g) from protein (20%), fat (10%), carbohydrate (70%)] until 8 weeks of age. At that time, mice were randomly assigned to either standard chow or high fat diet [TestDiet (St. Louis, MO) #58126: energy (kcal/g) from protein (18.3%), fat (60.9%), carbohydrate (20.1%)] for a further 8 weeks. For HFD cohorts, fed state body mass was assessed weekly from 8 weeks of age to 16 weeks of age between 11:00 and 13:00 hours. Blood glucose via tail-nick was also assessed at this time (OneTouch Ultra glucometer; maximum 34mM; LifeScan, Malvern, PA). At 0, 4, and 8 weeks of HFD, WT and Xin -/- mice were fasted for 6 hours and an intraperitoneal glucose tolerance test (IPGTT) was performed (1g/kg body weight at week 0 and 4; 0.5g/kg body weight at week 8). Blood glucose levels were assessed by tail-nick at 0, 20, 40, 60, 90, and 120 minutes.

### Tissue collection

Animals were euthanized via cervical dislocation at 16 weeks of age. The tibialis anterior (TA), gastrocnemius plantaris (GP) and soleus (Sol) muscles were rapidly harvested and coated in optimal cutting temperature (O.C.T.) compound (Leica Biosystems, Richmond, IL) before freezing in liquid nitrogen cooled 2-methylbutane. The quadriceps muscle group was flash frozen in liquid nitrogen. All samples were stored at −80°C until use.

### Cell Culture

Proliferating C2C12 cells (ATCC, CRL-1772) were cultured on polystyrene dishes (Fischer Scientific) in growth media containing high-glucose (4g/L) Dulbecco’s Modified Eagle Medium (DMEM) supplemented with sodium pyruvate and L-glutamine, 10% fetal bovine serum (FBS). Growth media was changed every 2 days. Cells were incubated at 37°C with 5% CO_2_ and kept at 25-80% confluence until passage number became prohibitively high.

### Adenoviral overexpression

To increase endogenous Xin expression, C2C12 cells were incubated with a custom recombinant designed XIRP-1 (Abm, Richmond BC; #C070-42417,) C-terminally tagged with human influenza hemagglutinin (HA). The adenoviral system is driven by a strong cytomegalovirus (CMV) promoter, replication incompetent due to E1/E3 deletions and human adenovirus type 5. Cells were washed once with warm sterile PBS and incubated with trypsin for 5-10mins at 37°C, 5% CO2. Using a hematocytometer each plate was seeded with 200,000 cells and given 2 hours to adhere. Following adherence, cells were infected for 6 hours. Viability of cells was accounted for by 0.4% Trypan Blue staining. For the duration of the infection, cells were kept in low volume of growth media (300uL) to promote virus incorporation. Control cells were infected with empty vector adenovirus (Vector Biolabs, Malvern PA; # 7028). After infection, cells were returned to normal confluence in growth media for 24 hours to allow for sufficient protein production.

### Mitotracker and Xin-HA staining

C2C12 myoblasts were seeded onto culture slides (BD Biosciences, Mississauga ON; #354104) and then infected with either ADV-HA-Xin or ADV-Null and incubated for 24 Hrs to allow for ample protein production. MitoTracker Red CMXRos (Invitrogen, Burlington ON; #M7512) and DAPI (Thermofisher, Mississauga ON; #D1306) were diluted in sterile growth media at a final concentration of 50nM and 10ug/mL respectively. Cells were incubated in the staining solution under normal culture conditions for 30mins, washed briefly in 1XPBS, and then visualized by epi- fluorescence microscopy (Nikon Canada, Mississauga ON; Eclipse Ti) for validation. Wells were washed 3 times for 5 mins in warm 1XPBS and later fixed in 4% paraformaldehyde for 7mins at room temperature. Wells were washed in cold 1XPBS (5 mins X3) and blocked for 1 hour at room temperature. Blocking reagent contained 5% normal goat serum (Vector Labs, Burlingame, CA, #S-1000), 0.3% Triton X-100 (Biorad, Mississauga ON; #1610407) diluted in 1X PBS. Primary antibody mouse α-HA tag- 1:100 (Cell Signaling Technology, Whitby ON; #6E2) diluted in antibody dilution buffer was added in enough volume to cover slides and incubated overnight at 4°C. Antibody dilution buffer contains 1% dissolved bovine serum albumin (Bioshop, Burlington ON; #Alb007.5) and 0.3% Triton X-100. Wells were washed in cold 1XPBS (5 mins X3) and then Alexa Fluor^TM^ 488 goat anti-mouse IgG1 (Thermofisher; #MG120) 1:2000 was added for 2 hours at room temperature in dark. Slides were briefly washed again in 1X PBS and then mounted in fluorescence mounting medium (Dako, Burlington ON; #S3023). Slides were dried overnight in the dark and then visualized using fluorescence microscopy (Nikon; Eclipse Ti).

### Histochemical and immunofluorescent stains

Frozen muscle samples (tibialis anterior) were freshly cut (6 um) and the enzymatic activity of Cytochrome Oxidase IV (COX; Complex IV) and succinate dehydrogenase (SDH) was assessed through (as described Shortreed et al., 2009). SDH and COX densities were calculated using threshold detection system of NIS-Elements software (Melville, NY, USA) and densities relative to fiber areas in muscle cross-sections were calculated.

Frozen longitudinal muscle sections muscle sections (tibialis anterior) were also co-stained with anti-Xin antibody (Hawke et al., 2007, Nilsson et al., 2013, Al-Sajee et al., 2015) and anti-Tom20 or anti-Desmin, antibodies, along with the appropriate secondary antibodies to visualize the co-localization of Xin with Tom20 or Desmin respectively. Images were acquired using Nikon 90i eclipse upright microscope and Olympus (Richmond Hill., ON; FV1200) laser scanning confocal microscope.

### Mitochondrial respiration

#### Permeabilized Muscle Fibres

Permeabilized muscle fibres were prepared as previously described (Monaco et al., 2018). Briefly, the red TA muscle from the right leg was harvested from WT and Xin-/- mice. Muscle was immediately placed in ice-cold BIOPS buffer containing 50mM MES, 7.23mM K_2_EGTA, 2.77mM CaK_2_EGTA, 20mM imidazole, 0.5mM dithiothreitol (DTT), 20mM taurine, 5.77mM ATP, 15mM PCr, 6.56mM MgCl_2_×6H_2_O (pH 7.1). The muscle was then separated into several small muscle bundles and each bundle was gently separated along the longitudinal axis using needle-tipped forceps. Following this, bundles were treated with 40μg/mL saponin and 1μM CDNB in BIOPS buffer on a rotor at 4°C for 30 minutes to allow for permeabilization of the sarcolemma and depletion of glutathione as previously described (Fisher-Wellman et al., 2013). Following incubation, fibres were washed in Buffer Z containing 105mM K-MES, 30mM KCl, 10mM KH_2_PO_4_, 5 MgCl_2_×6H_2_O, 1mM EGTA, 5mg/mL BSA (pH 7.4) at 4°C.

#### Mitochondrial Respiration

The Oroboros O2k (Oroboros Instruments, Corp., Innsbruck, Austria) system was used to measure respiration in permeabilized muscle fibres at 37°C with constant magnetic stirring at 750rpm. Permeabilized fibres were added to 2mL of Buffer Z respiration medium containing 20mM creatine, 20 µM Amplex Red, 5 U/ml horseradish peroxidase, and 40 U/ml superoxide dismutase 1. Chambers were hyper-oxygenated to 350-375μM. State IV respiration was measured in the presence of 5mM pyruvate and 2mM malate. and chambers were hyper-oxygenated to 350-375μM. State IV respiration was measured in the presence of 5mM pyruvate and 2mM malate. State III respiration was then measured by addition of 5mM ADP. Mitochondrial outer membrane integrity was next verified by adding 10μM cytochrome *c* and samples with a >10% increase in respiration were excluded. For complex II specific respiration, complex I was then inhibited with 0.5μM rotenone followed by 10mM succinate in the same bundles. *J*O_2_ was normalized by dividing by fiber bundle wet weight as well as normalizing to bundle wet weight and mitochondrial density (as assessed by electron microscopy). Unfortunately, analysis of mitochondrial H_2_O_2_ emission was not possible as values were below the level of detection for reasons unknown. For consistency between all sample analyses however, we remained consistent in the addition of all reagents and buffers.

### Electron microscopy imaging and analysis

Left tibialis anterior muscles were collected from WT and Xin-/- mice and fixed in 4% paraformaldehyde with 1.5% glutaraldehyde. Longitudinal sections were prepared on copper grids, contrasted with uranyl acetate and lead citrate and viewed in a JEOL JEM 1200 EX TEMSCAN transmission electron microscope (JEOL, Peabody, MA) at McMaster University. Images were captured from non-overlapping sections at 30,000x magnification. 30,000x magnification images were subsequently analyzed using Nikon NIS-Elements ND2 software. Mitochondrial area (total mitochondrial area), density (total mitochondria area/total fibre area, reported as a percentage of the total fibre area), content (number of individual mitochondria per total fibre area), mean electron density (mitochondrial electron-density) were measured via manual circling of individual mitochondria. The distance between Ryanodine receptors (RyR) side of SR and outer mitochondrial membrane was measured using 30000X magnification with a total of 60-80 SR and outer mitochondrial membrane included as described in (Boncompagni et al., 2009). This distance represents the presumed pathway of Ca^2+^ to travel from SR to mitochondria.

### Immuno-electron microscopy (ImmunoEM)

Surgical discard skeletal muscle samples from knee and hip arthroplasty patients were fixed with 4% paraformaldehyde and 1.5% glutaraldehyde within 10 minutes of removal and embedded in LR-white resins. These ultrathin sections were prepared and blocking reagent consisting of 10% bovine serum albumin in filtered phosphate buffered saline (PBS) was applied for 10 minutes.

Ultrathin sections were then incubated with anti-Xin antibody (in 10% BSA) for 1 hour at room temperature. Goat Anti-Chicken Gold 6nm particles were used as a secondary antibody. Uranyl acetate and lead citrate were used as counter stains. Silver enhancement was not used. Images were acquired with an AMT 4-megapixel digital camera (Advanced Microscopy Techniques, Woburn, MA). All procedures were approved by the McMaster University Human Integrated Research Ethics Board (HiREB #1614) and conformed to the Declaration of Helsinki.

### Western blot analysis

Western blotting was completed using standard procedures with snap-frozen quadriceps muscle from WT and Xin-/- mice. Muscle was homogenized in lysis buffer and protein content was determined using a BCA assay. Lysates were run on 10, 12%, or 15% SDS-PAGE gel, depending on the molecular weight of the protein being investigated, and then transferred to PVDF membrane, with the exception of Xin lysates which were transferred to a nitrocellulose membrane. Primary antibodies (p62, Drp1, Mfn1, Mfn2, pULK, PGC1a, PFK1, LDH, HKII, GAPDH, Opa1) were incubated overnight at 4°C. Appropriate horseradish conjugated secondary antibodies were applied. Chemiluminescent reagent (Biorad #1705061) was used to visualize proteins of interest. All information pertaining to antibodies is provided in Table 1.

**Table 1.** Antibody information.

| Antibody | dilution | Company/Catalogue # |
| --- | --- | --- |
| Mouse monoclonal primary antibodies |  |  |
| Xin | 1:5 (1:500 for EM) | HybroTec #XR1A |
| Opa1 | 1:1000 | BD Transduction; #612606 |
| Drp1 | 1:1000 | Abcam; #56788 |
| Mfn1 | 1:1000 | Abcam #57602 |
| Mfn2 | 1:1000 | Abcam #56889 |
| Desmin | 1:150 | Vector Laboratories; #VP-D502 |
| Tom20 | 1:100 | Santa Cruz; #sc-17764 |
| HA-tag | 1:100 | Cell Signaling; #6E2 |
| Rabbit polyclonal primary antibodies |  |  |
| pULK1 | 1:1000 | Cell Signalling #5869 |
| PGC1a | 1:1000 | Santa Cruz; #13067 |
| LDH | 1:1000 | Cell Signaling; # 3558s |
| PFK-1 | 1:1000 | Abgent; #AP8137a |
| p62 | 1:1000 | Cell Signaling; #8025 |
| Hexokinase II | 1:1000 | Cell Signaling; #2867 |
| GAPDH | 1:2000 | Thermo (Pierce); # PAI-987 |
| Secondary Antibodies |  |  |
| Goat anti-Rabbit IgG H&L (HRP) | 1:10000 | Abcam; #6721 |
| Goat anti-Mouse IgG H&L (HRP) | 1:10000 | Abcam #6789 |
| Goat Anti-Mouse IgG H&L (Alexa Fluor® 488) | 1:1000 | Abcam; #150113 |
| Goat Anti-Mouse IgG H&L (Alexa Fluor® 594) | 1:1000 | Abcam; #150116 |
| Goat Anti-Chicken IgY H&L Gold 6nm | 1:30 | Abcam; #39604 |

### Statistical analysis

The N values for each experiment are provided within the figure legends. All statistical analysis was conducted using GraphPad Prism 7.0 (San Diego, CA). As appropriate, a Student’s t-test or two-way ANOVA was used for comparison between groups and Bonferroni post-hoc analysis was performed as needed. Statistical significance was set at a p value of less than 0.05. All data presented are mean ± SEM (standard error of the mean).

## Results

### Xin protein is present in the mitochondrial/peri-mitochondrial regions of skeletal muscle

Using adenoviral infection to overexpress an HA-tagged Xin in C2C12 myoblasts, a clear co-localization was observed between HA-tagged Xin and Mitotracker™ (fluorescent dye that stains mitochondria in live cells; Pearson’s coefficient of 0.61) (Fig 1A). This observation was further supported through laser scanning confocal microscopy analysis of human skeletal muscle sections which demonstrated significant overlap in Xin expression with the mitochondrial-specific protein, Tom20 (Fig 1B; Pearson’s coefficient of 0.46). Furthermore, immuno-EM found focal aggregates of electron dense particles [anti-Xin antibody with Goat Anti-Chicken Gold secondary] in the mitochondrial and peri-mitochondrial regions of human skeletal muscle (Fig 1C). Collectively, these findings support the presence of Xin in and around the mitochondria of murine and human skeletal muscle.

**Figure 1:**
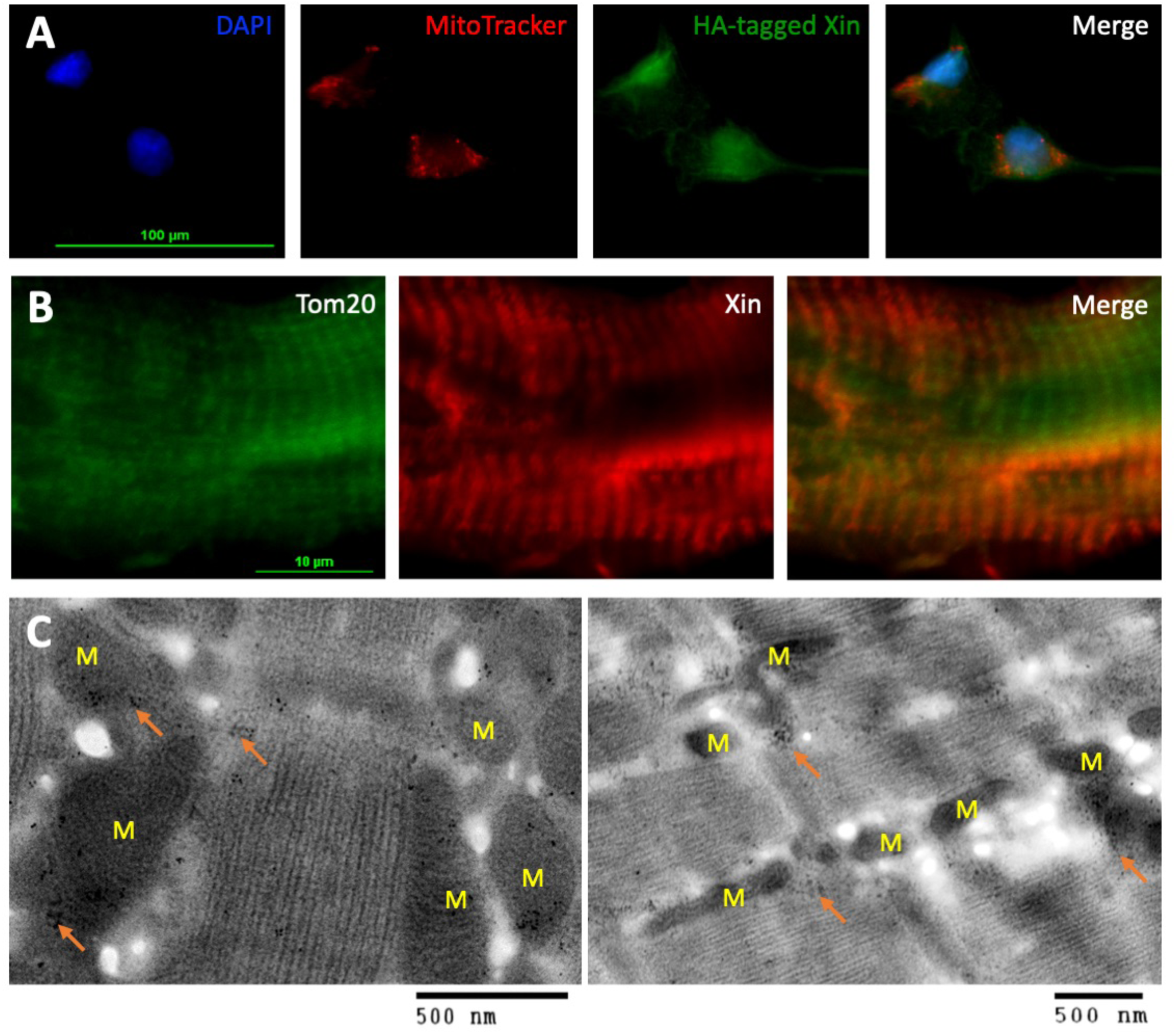
Xin expression in skeletal muscle consistent with mitochondrial and peri- mitochondrial regions. (A) Adenoviral infection of HA-tagged Xin in C2C12 myoblasts allowed for a clear co-localization between HA-tagged Xin and Mitotracker™ (Pearson’s coefficient of 0.61). (B) Confocal microscopy analysis of human skeletal muscle samples (longitudinal sections) demonstrated significant overlap in Xin expression with the mitochondrial-specific protein Tom20. A Pearson’s coefficient of 0.46 was recorded. (C) Immuno-electron microscopy of paraformaldehyde-fixed skeletal muscles revealed focal aggregates of electron dense particles [anti-Xin antibody with Goat Anti-Chicken Gold secondary] in the mitochondrial and peri-mitochondrial regions of human skeletal muscle (orange arrows denote electron dense particle clusters in the areas within and surrounding the mitochondria (denoted by an M). Collectively, these findings point to the presence of Xin in and around the mitochondria of murine and human skeletal muscle.

### Skeletal muscles of mice lacking Xin display abnormal mitochondrial morphology but increased mitochondrial enzyme content

We next interrogated the effect of Xin deficiency on mitochondrial morphology. While WT intermyofibrillar (IMF) mitochondria were consistently observed as small, dark grey (electron dense) circles located on each side of the Z-line (Fig 2A,B), the IMF mitochondria in Xin-/- muscles were frequently forming elongated networks often spanning the area between two Z-lines with a significant number exhibiting swelling (reduced electron density) and abnormal cristae patterning (Fig 2C,D). What became visible on the ultrastructural images were the notable increases in mitochondrial content, specifically IMF (but not sub-sarcolemmal) mitochondria. Skeletal muscle cross-sections were used to assess succinate dehydrogenase (SDH) enzyme activity as a marker of mitochondrial proliferation. Confirming the electron microscopy observations, we measured a significant increase in SDH enzyme activity staining in Xin-/- muscles without the thickened ring of SDH staining which characterizes ragged blue fiber staining (Fig 2E-G). We then undertook Cytochrome Oxidase IV (COX) staining, to investigate whether there were myofibers which were COX negative; used to visualize impairments to mitochondrial respiratory chain (Fig 2H-J). We found no differences between cohorts with respect to the presence of COX-negative myofibers indicating the existence of increased mitochondrial content in the absence of overt mitochondrial disease.

**Figure 2.**
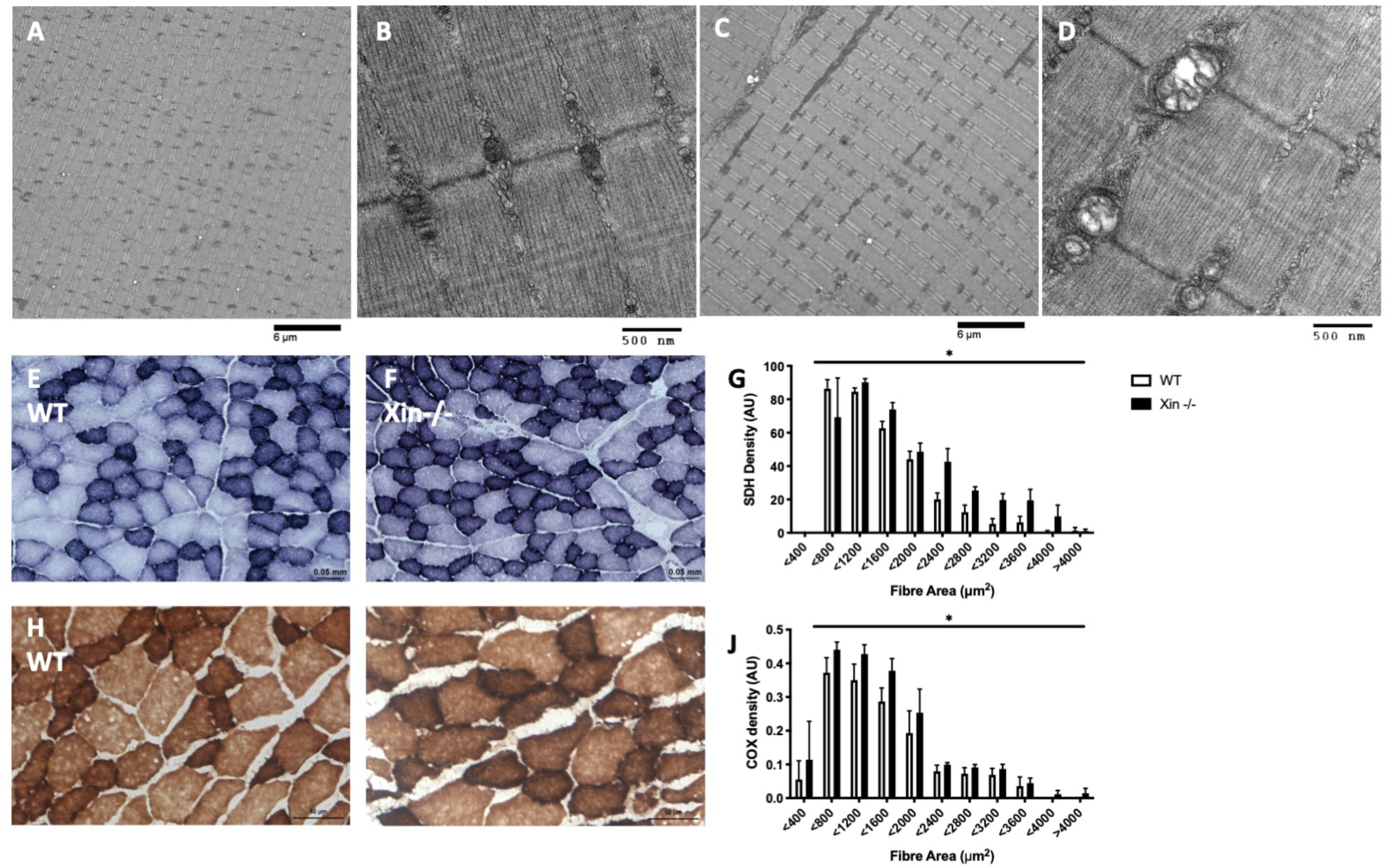
Alterations to mitochondrial morphology and increased mitochondrial enzyme activities in Xin deficient mouse muscle. (A & B) Intermyofibrillar (IMF) mitochondria were consistently observed as small, dark grey (electron dense) circles located on each side of the Z-line in Wildtype (WT mouse skeletal muscle, whereas (C&D) the IMF mitochondria in Xin-/- muscles were frequently forming elongated networks often spanning the area between two Z-lines with a significant number exhibiting swelling (reduced electron density) and abnormal cristae patterning. (E-G) Skeletal muscle cross- sections were then used to assess succinate dehydrogenase (SDH) enzyme activity as a marker of mitochondrial content. Confirming the electron microscopy observations, we measured a significant increase in SDH enzyme activity staining in Xin-/- muscles without the thickened ring of SDH staining which characterizes ragged blue fiber staining (see E&F). We also undertook Cytochrome Oxidase IV (COX) enzymatic staining (H-J), to investigate whether myofibers were COX negative. COX negative fibers are used to visualize impairments to mitochondrial respiratory chain and is a hallmark sign of mitochondrial myopathy. We found no differences between cohorts with respect to the presence of COX negative myofibers though we did see a generalized increase in COX staining density (consistent with SDH staining); indicating the existence of increased mitochondrial content in the absence of overt mitochondrial disease. * represents a main effect between groups of p<0.05 as assessed by a 2 Way ANOVA with n=4 per group for all measures with greater than 75 myofibers per muscle analyzed.

### No difference in weight gain and blood glucose handling between WT and Xin-/- mice in response to HFD

Given the significant increases in mitochondrial density and alterations to mitochondrial morphology in resting, chow-fed Xin-/- skeletal muscles, we hypothesized that providing a metabolic stress in the form of a high fat diet for 8 weeks would elicit significant myopathic changes in these mice.

As expected, significant weight gain was observed in both groups, surprisingly however, no significant differences were observed between the control and Xin-/- mice throughout the experimental period (Fig 3A). Furthermore, no differences in glucose handling were observed between WT and Xin-/- mice on a HFD as assessed by fasting blood glucose levels (Fig 3B), and intraperitoneal glucose tolerance tests at 4 weeks (Fig 3C-F) and 8 weeks (Fig 3G-J) of a HFD.

**Figure 3.**
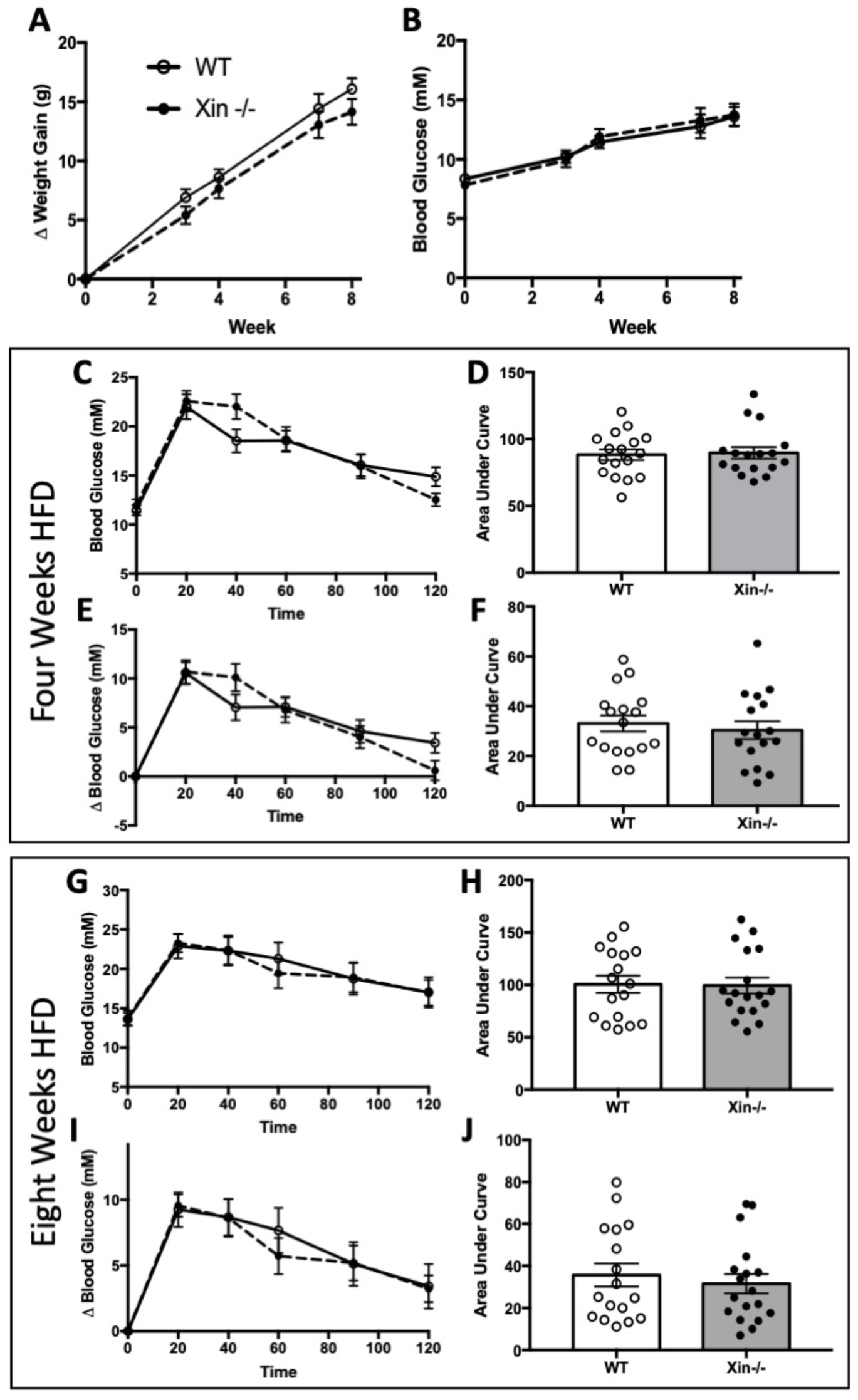
No difference between groups in weight gain or blood glucose control with the feeding of a high fat diet. (A) Significant weight gain was observed in both groups but no significant differences between groups were observed throughout the experimental period (B) No differences in fasting blood glucose was observed between WT and Xin-/- mice fed a HFD. Intraperitoneal glucose tolerance tests at four weeks (C-F) and eight weeks (G-J) following HFD initiation also demonstrated no differences between groups in glucose tolerance. N=17 per group.

### Skeletal muscle mitochondria in high-fat fed Xin-/- mice display abnormal morphology, increased density, and reduced respiration capacity

To assess mitochondria morphology and content, murine tibialis anterior muscles were analyzed using electron microscopy. Intermyofibrillar mitochondria within WT mice appeared small and localized peripherally to the Z-discs with limited abnormalities (Fig 4A). In contrast, the mitochondria of Xin-/- skeletal muscle displayed extensive swelling and streaming, along with disruption of cristae (Figure 4B). Quantitatively, the average IMF mitochondria of Xin-/- skeletal muscle displayed significant increase in area (Fig 4C) with a significant reduction in cristae density [as assessed by mean electron density of each mitochondria (Fig 4D)]. Given the significant increase in mitochondrial area, it is not surprising there was also a significant increase in mitochondrial density in Xin-/- muscles (Fig 4E). In contrast to IMF mitochondria, SS mitochondria in Xin-/- mice did not exhibit a significant change in mitochondrial density relative to WT littermates (data not shown).

**Figure 4.**
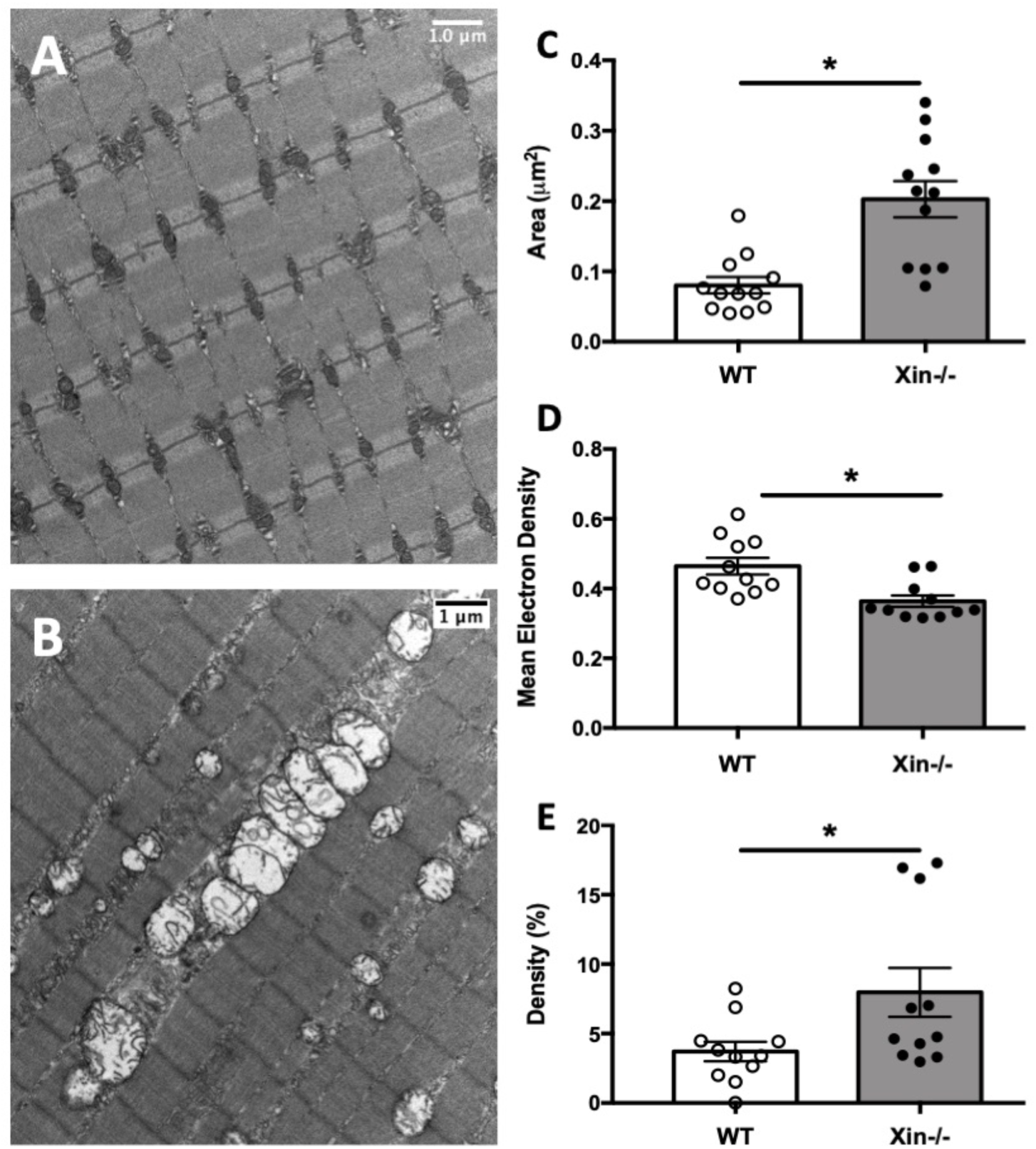
Xin-/- skeletal muscles exhibit increased mitochondrial area, density, as well as reduced electron density. Following HFD feeding, intermyofibrillar mitochondria within WT mice appeared small and localized peripherally to the Z-discs with limited abnormalities (A). In contrast, the mitochondria of Xin-/- skeletal muscles displayed extensive swelling and streaming, along with disruption of cristae (B). Quantitatively, individual IMF mitochondria of Xin-/- skeletal muscle displayed significant increases in (C) area with a significant reduction in (D) cristae density, assessed by mean electron density of each mitochondria. Given the significant increase in mitochondrial area, it is not surprising there was also (E) a significant increase in overall IMF mitochondrial density in Xin-/- muscles. * represents a difference between groups of p<0.05 with n=11 per group.

To determine if these morphological alterations were manifesting as an impairment in mitochondrial function, permeabilized muscle fibres from WT and Xin-/- mice were analyzed using high resolution respirometry. Despite the significant abnormalities in mitochondrial morphology in Xin-/- skeletal muscles, no significant difference in maximal complex I respiration (Fig 5A) or complex II respiration (Fig 5B) when calculated per milligram of wet muscle. These results would suggest that the elevated mitochondrial density in Xin-/- muscles may be a compensatory mechanism. To estimate mitochondrial respiration on a per individual mitochondria level, we made mitochondrial respiration values relative to mitochondrial density. Our hypothesis was supported by significant decreases in both complex I (Fig 5C) and II (Fig 5D)-supported respiration/mitochondrial density in Xin-/- skeletal muscle.

**Figure 5.**
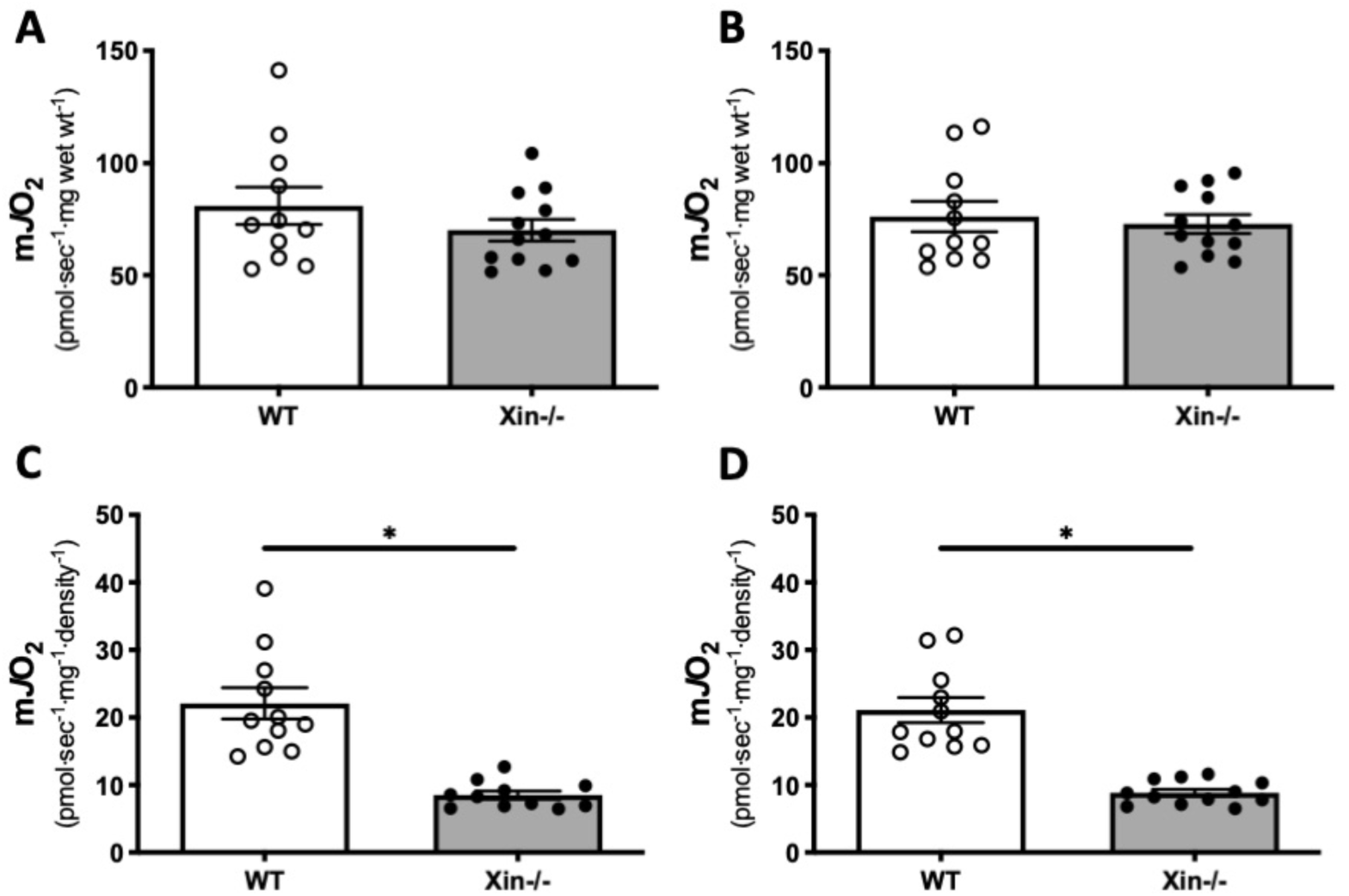
High resolution respirometry measurements revealed impaired mitochondrial function at the mitochondrial level which was compensated for by increased mitochondrial density. Permeabilized muscle fibres from WT and Xin-/- mice were analyzed using high resolution respirometry. No significant differences in maximal complex I (A) or complex II (B) respiration were observed between groups (per milligram of muscle wet weight). However, when the data in (A) and (B) were normalized to IMF mitochondrial density (per mg wet weight/mito density), significant decreases in both complex I (C) and complex II (D) -supported respiration per unit density in Xin-/- skeletal muscle were observed. * represents a difference between groups of p<0.05 with an n =11 for WT and n=12 for Xin-/-.

### No change in mitochondrial dynamics proteins or in key glycolytic enzymes

In order to ascertain if the changes to the mitochondrial structure and function were the result of adaptations to mitochondrial fission, fusion, or autophagy, we used immunoblotting to assess the levels of key mitochondrial dynamics proteins. Changes to mitochondrial dynamics proteins could help explain the morphologically-abnormal mitochondria observed in Xin-/- mice. However, immunoblot analysis of mitochondrial fusion (MF1, MF2, OPA1), fission (DRP1), and autophagy (pULK, p62) proteins revealed no differences between WT and HFD mice in any metric (Fig 6A- F). We also observed no significant difference in the protein expression of PGC-1a (Fig 6G) or key glycolytic enzymes: lactate dehydrogenase (LDH; Fig 6H), phosphofructokinase (PFK; Fig 6I) and hexokinase II (HKII; Fig 6J) between WT and Xin-/- mice on a HFD suggesting that these alterations in mitochondria were not resulting in a change in glycolytic enzyme expression.

**Figure 6.**
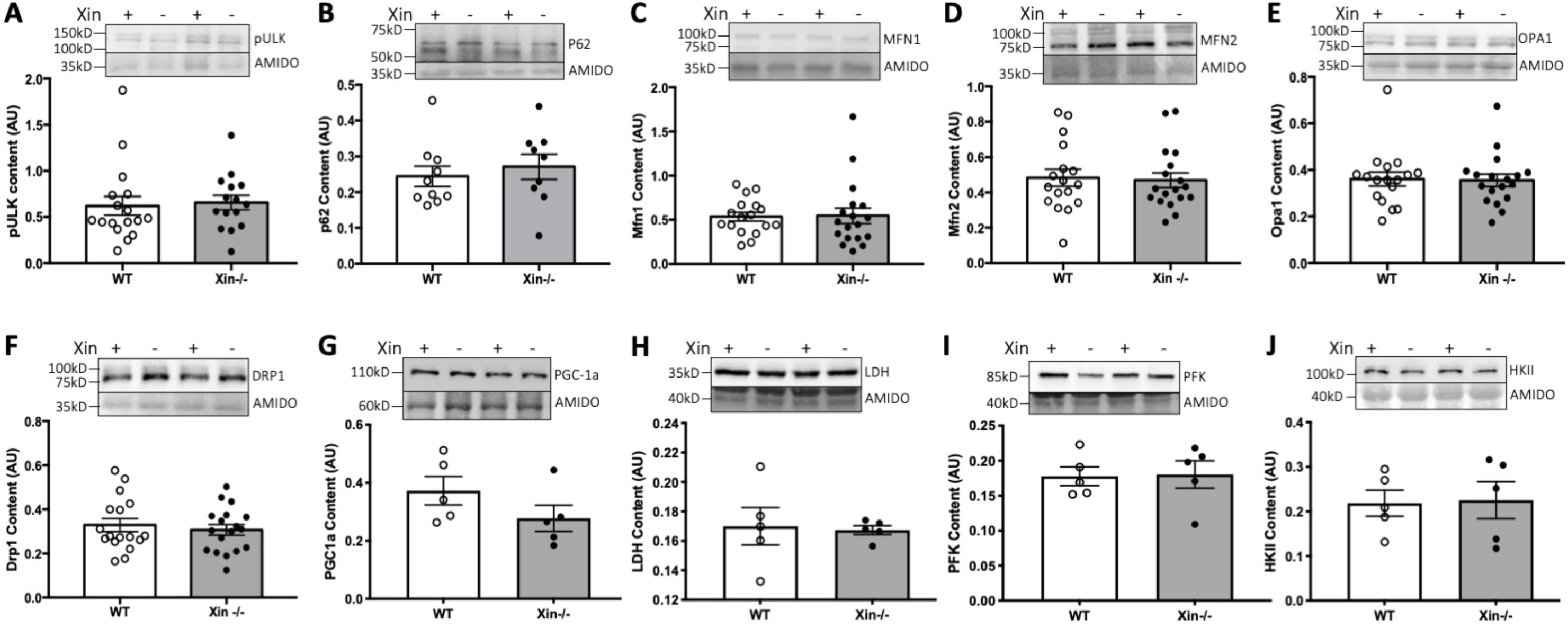
Immunoblotting of key mitochondrial dynamics and glycolysis proteins. Immunoblot analysis of mitochondrial autophagy [(A) pULK, (B) p62], fusion [(C) Mfn1, (D) Mfn2 (E) Opa1] and fission [(F) Drp1] proteins revealed no differences between WT (Xin+) and Xin-/- (Xin-) mouse muscle in any metric. In addition, no significant differences in the protein expression of (G) PGC-1a or key glycolytic enzymes (H) lactate dehydrogenase (LDH), (I) phosphofructokinase (PFK) and (J) hexokinase II were observed between WT and Xin-/- mice. N for each group is highlighted by individual datapoints for each graph bar.

### Xin co-localizes with the intermediate filament protein, Desmin, and the absence of Xin alters intermyofibrillar mitochondria ultrastructural location

Alterations to the morphology of intermyofibrillar mitochondria in Xin-/- skeletal muscles without a notable change in mitochondrial dynamics proteins would suggest that Xin may be involved in the sarcoplasmic reticulum (SR)- mitochondria interface. The mitochondrial-associated membranes (MAMs) are a critical interface with known roles in Ca^2+^ and lipid transfer between SR and mitochondria. However, recent work has also shown a key role for MAMs in the regulation of mitochondrial shape and motility, energy metabolism and redox status (van Vliet et al., 2014). Here we show that Xin colocalizes with the intermediate filament protein, and known MAM protein, Desmin (Milner et al., 2000) with a Pearson’s coefficient of 0.72 (Fig 7A). In addition to the change in ultrastructural morphology in mitochondria from Xin-/- muscle, we also evaluated the relationship of IMF mitochondria to the triads. The distance from SR membrane (RyR1) to the outer mitochondrial membrane, which is the presumed path for Ca^2+^ to travel from the release channels to mitochondria, was significantly smaller in Xin-/- compared to WT muscles (Fig 7B,C).

**Figure 7.**
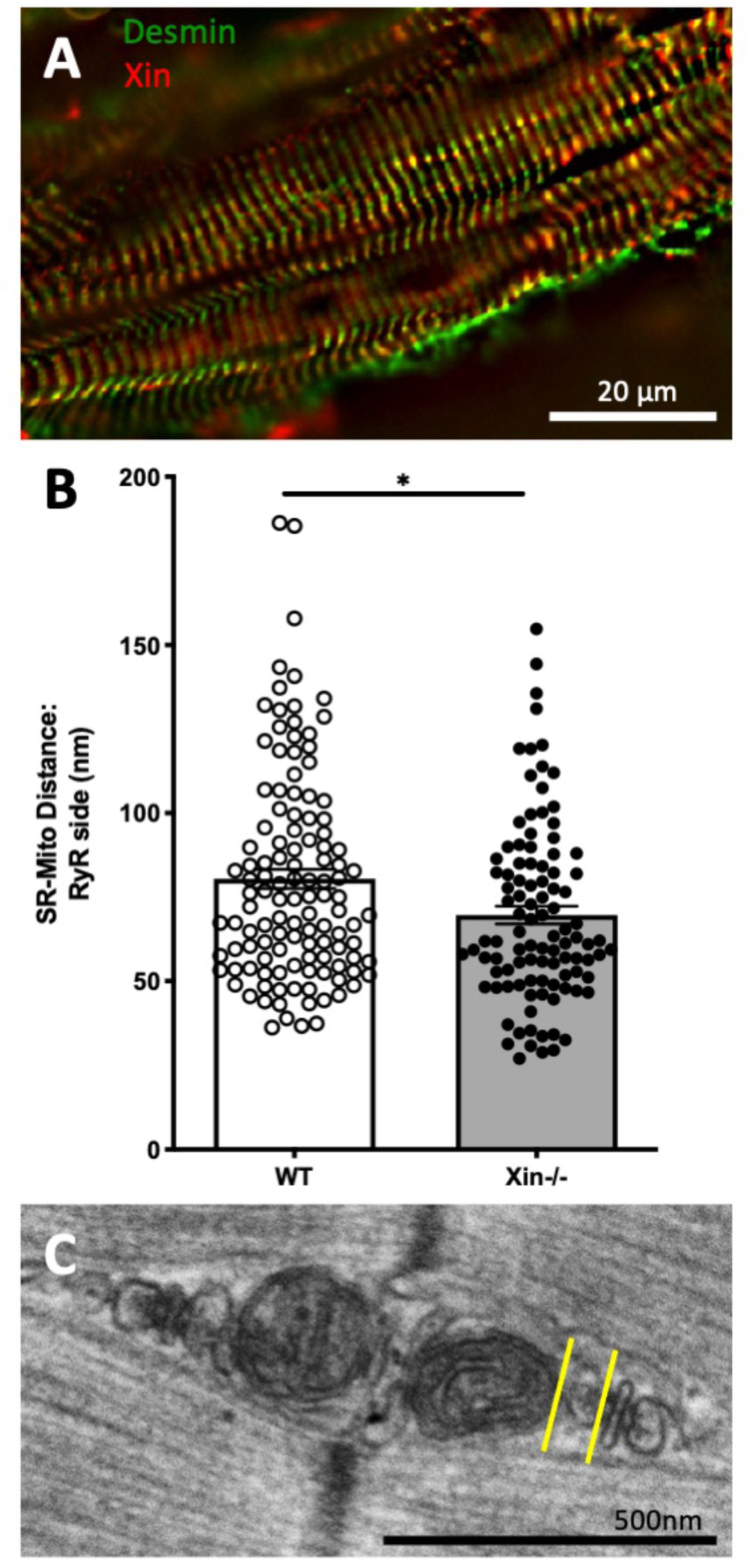
Xin co-localizes with Desmin and the loss of Xin alters SR-mitochondrial distance. Xin colocalizes with the intermediate filament protein, and known MAM protein, Desmin with a Pearson’s coefficient of 0.72 (A). In addition to the change in ultrastructural morphology in mitochondria from Xin-/- muscle, we also evaluated the relationship of IMF mitochondria to the triads. The distance from SR membrane (RyR1) to the outer mitochondrial membrane, which is the presumed path for Ca^2+^ to travel from the release channels to mitochondria, was significantly smaller in Xin-/- compared to WT muscles (B, illustrated in panel C as the distance between the yellow lines).

## Discussion

The interplay between cytoskeletal proteins and mitochondria in skeletal muscle is an area of intense interest and a number of important players are coming to light. Here, we provide compelling evidence that Xin, a cytoskeletal adapter protein, may be involved in this interface and the absence of Xin alters IMF mitochondrial density and localization.

The presence of Xin in the mitochondrial/peri-mitochondrial regions of skeletal muscle has not been reported in these areas previously with the predominance of literature suggesting that this multi-adapter protein is localized specifically to sites with considerable mechanical strain, such as the myotendonous junctions and the costameres of skeletal muscle, and intercalated discs of the heart (VanderVen 2006; Dev Dyn. 2002; 225: 1-13; Al-Sajee 2015; Hawke et al. 2007; Nissar- doi: 10.1152/ajpcell.00298.2011). The presence of Xin in (peri)-mitochondrial regions of muscle appears to be more than a casual position however, as the absence of Xin results in a significant perturbation in mitochondrial morphology and an altered ultrastructural distribution. Intermyofibrillar mitochondria are dependent on numerous cytoskeletal connections for proper localization and function within skeletal muscle (Anesti et al). The loss of this mitochondria- cytoskeletal connectivity can significantly impact mitochondrial morphology and function as has been reported for a number of other structural proteins such as Desmin (Milner et al., 2000; Carlsson et al., 2002; Paulin et al., 2004; Schwartz et al., 2016), Cav-3 (Shah et al., 2020) and cytoplasmic actin isoforms (βcyto- and γcyto-actin; O’Rourke et al., 2018). Loss of any of these proteins has been reported to result in morphological alterations to skeletal muscle mitochondria; similar in many aspects to what is observed with Xin deficiency herein. While Xin has not been reported to have direct interaction with Desmin or either cytoplasmic actin isoforms, we did observe a measurable overlap of Desmin and Xin staining in skeletal muscle that indicates proximity of these two proteins, likely to the MAM. It is also worth noting that a query of Xirp1 within the String™ Interaction Network (Szklarczyk et al., 2019) indicates that Xin is predicted to interact with Caveolin-3 (Cav3) with high confidence, and it is possible that this relationship may be disrupted leading to mis-localization or mis-expression of Cav3, and ultimately, be a contributor, along with Desmin, to the observed changes in mitochondrial morphology. Clearly future studies are warranted to detail the potential relationships between Xin, Desmin and Cav3 within skeletal muscle.

The proliferation of mitochondria in Xin-/- skeletal muscle appears to be a compensatory mechanism allowing for the maintenance of mitochondrial respiration even in the face of mitochondrial swelling and loss of cristae density. Furthermore, despite these significant abnormalities in mitochondrial morphology and localization, the challenge of a high-fat diet did not elicit differences in body weight or blood glucose handling in Xin-/- mice compared to their WT littermates. The lack of difference in the measured mitochondrial dynamics proteins and key glycolytic enzymes suggest that the alterations in mitochondrial morphology resulting from the absence of Xin further support the hypothesis of a role for Xin in cytoskeleton-mitochondrial interaction rather than a role in mitochondrial dynamics. We do acknowledge however that protein expression changes do not always reflect a dynamic process such as fission, fusion or mitophagy and thus more specific investigations are warranted to rule out a role for Xin in regulating mitochondrial dynamics.

In summary, the present results identify a heretofore unrecognized location and role for the cytoskeletal adapter protein, Xin, in skeletal muscle. Our findings provide evidence for Xin being localized in the (peri)-mitochondrial regions, with a possible involvement in the mitochondrial-associated membranes; such that the absence of Xin results in significant abnormalities in mitochondrial morphology without notable detriment to mitochondrial dynamics.

## Contribution statement

All authors provided final approval of the version to be published. All people designated as authors qualify for authorship, and all those who qualify for authorship are listed. TJH is the guarantor of this work and, as such, had full access to all the data in the study and take responsibility for the integrity of data and the accuracy of the data analysis.

